# Predicting “pain genes”: multi-modal data integration using probabilistic classifiers and interaction networks

**DOI:** 10.1101/2024.05.15.594305

**Authors:** Na Zhao, David L Bennett, Georgios Baskozos, Allison M Barry

## Abstract

Accurate identification of pain-related genes remains challenging due to the complex nature of pain pathophysiology and the subjective nature of pain reporting in humans, or inferring pain states in animals on the basis of behaviour. Here, we use a machine learning approach to identify possible “pain genes”. Labelling was based on a gold-standard list of genes with validated involvement across pain conditions, and was trained on a selection of -omics (eg. transcriptomics, proteomics, etc.), protein-protein interaction (PPI) network features, and biological function readouts for each gene. Multiple classifiers were trained, and the top-performing model was selected to predict a “pain score” per gene. The top ranked genes were then validated against pain-related human SNPs to validate against human genetics studies. Functional analysis revealed JAK2/STAT3 signal, ErbB, and Rap1 signalling pathways as promising targets for further exploration, while network topological features contribute significantly to the identification of “pain” genes. As such, a PPI network based on top-ranked genes was constructed to reveal previously uncharacterised pain-related genes including CHRFAM7A and UNC79. These analyses can be further explored using the linked open-source database at https://livedataoxford.shinyapps.io/drg-directory/, which is accompanied by a freely accessible code template and user guide for wider adoption across disciplines. Together, the novel insights into pain pathogenesis can indicate promising directions for future experimental research.

## Introduction

The identification of pain-related genes remains challenging due to the heterogeneity and multifactorial nature of the disease. “Pain” encompasses an unpleasant sensory and emotional experience, acute and chronic states, and actual or potential tissue damage spanning a spectrum of conditions from UV radiation (sunburn) to diabetic neuropathy (***Raja et al. (2020)***).

High-throughput sequencing techniques (“-omics”), have revolutionised the identification of molecular markers and pathways, with technologies like transcriptomics and translatomics providing large gene expression data sets which enable identification of differentially expressed genes and data-driven biomarker discovery. With such diversity in pain characterizations, we expect equally diverse underlying mechanisms. Even so, informative parallels are also seen across conditions. For example, decades of work in the migraine field have culminated to the development of clinical CGRP antibodies – a hallmark treatment for acute and chronic migraine (***Edvinsson et al. (2018); Goadsby et al. (2017)***). This pathway is also being explored in the context of neuropathic and inflammatory pain, as well as non-migraine headache pain due to the underlying mechanism of sensory neuron sensitization (***Schou et al. (2017); Paige et al. (2022)***).

As we continue to accumulate -omics data, we need effective strategies to present and integrate these datasets together, while also considering multi-modal data from external sources.

Machine learning (ML) approaches are well suited to this challenge, and the availability of probabilistic models (that is – models which give the probability something belongs to a specific class) allows us to assign a probability score to each instance. Recently, ML integration with gene expression data has gained popularity in biomarker discovery (reviewed in ***Zhang et al. (2021)***) and clinical diagnosis (***Kumar et al. (2023)***).

In the context of pain, this has proven to be powerful: for example, predicting patient classification for painful vs painless diabetic neuropathy has highlighted factors relevant to the painful class (***Baskozos et al. (2022)***). Pre-clinically, there is also a need to integrate multi-modal data, both to give insight to features underlying genes involved in pain, as well as to lay foundation for future studies.

Here, we trained both off-the-shelf probabilistic classifiers and ensembles of classifiers to produce a predictive pain score based on an expansive feature space to address this gap. Features include cross-species transcriptomic and translatomics datasets as well as proteomic data, network topology, genetic structure (eg. GC content), and functional pathway assignments.

The top-performing model was employed to predict a class probability score (“pain score”) for each gene, and the gene candidates with highly ranked predicted pain scores are subjected to downstream functional analysis, including validation against human genetics datasets. High ranking features, such as DRG translatomics data and protein-protein network interactions were highlighted and further examined in the context of pain, while JAK2/STAT3 signal, ErbB, and Rap1 signalling pathways were identified as promising targets for future exploration.

These scores were curated into an open-access database (https://livedataoxford.shinyapps.io/drg-directory/) alongside experimental datasets. Here, we have integrated the STRING DB to facilitate the visualisation of pain-related genes in the context of their network associations (a high ranked feature), building on previous work by Perkins et al., 2013 (***Perkins et al. (2013)***).

In addition to data integration, a fundamental component to effective data use is access. As more big data continues to be produced, better data management practices are needed (***Boeckhout et al. (2018)***): Raw data repositories limit use to those with bioinformatic skills while extensive supplemental data tables quickly become cumbersome.

We have thus paired our database to a user-guide for researchers to reimplement these visualization for other omics studies and disease states: This reproducible Shiny-based framework can simply integrate multiple omics datasets and generate composite visualisations and, where relevant, add PPI networks of condition-related genes to help inform candidate selection for downstream experimentation. Additionally, it addresses a legacy gap commonly seen with in-house databases.

Together, this study presents a novel disease-gene identification framework by integrating diverse datasets through machine learning to gain mechanistic insights of pain. Paired to an open access database with an emphasis on PPI networks, this will allow researchers to more effectively select targets, and, ultimately - lead to better data utilisation and increased impact of each study.

## Results

Here, we use a machine learning approach to identify possible “pain genes” (Figure 1). Because of the diversity in the studies used to generate initial labels, the term “pain” here is used as a broad characterization across acute and chronic states. Here, the predicted pain genes thus represent more generalisable genes across conditions, opposed to, for example, predictions tailored to neuropathic pain specifically.

**Figure 1.**
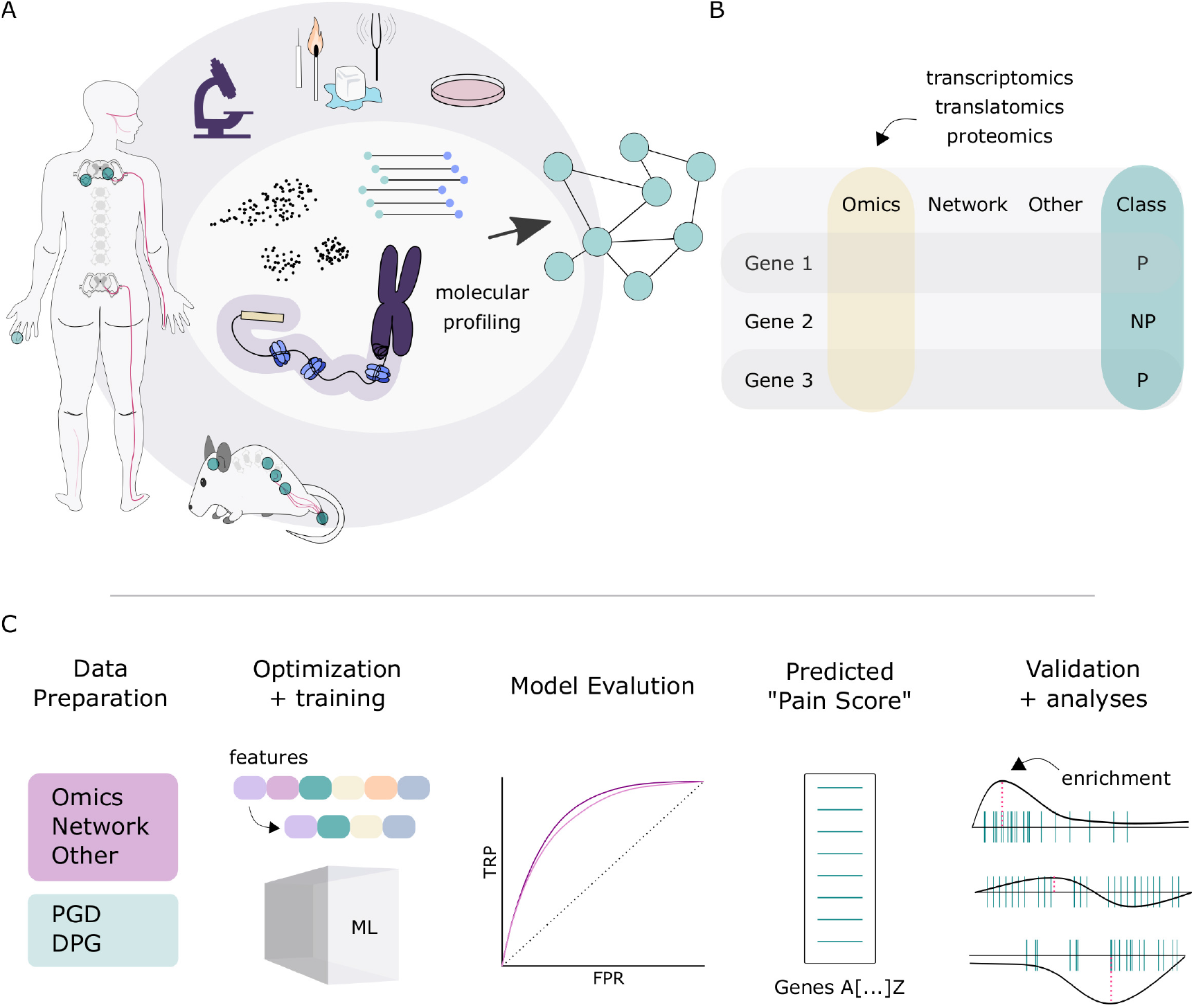
Experimental overview. A. Data integration schematic, where data can be integrated across species and modalities using machine learning. B Example data input, when information for each gene was gathered across modalities and labelled either pain (“P”) or no-pain (“NP”) based on gold-standard lists from prior literature. C. Pipeline schematic for predicting “pain” genes, from data preparation through to validation.

Selecting gene labels in the experimental design is not a trivial task, due to the variable amount of research surrounding each gene in the context for pain. Here, we opt for a highly stringent approach, requiring functional, in vivo validation in mice or detailed characterization in humans (see methods for full details). With this, there is an underlying expectation that a number of genes being studied in the context of pain do not yet reach our threshold for inclusion as “pain”, even though they are likely to be relevant: ie. gene classified as pain when labelled non-pain is not necessarily incorrect biologically, but a comment on our current knowledge in the field.

### Feature Selection and Exploration

Scikit-learn (Sklearn) was used to classify “pain”/”no pain” genes in Python using a variety of algorithms (***Pedregosa et al. (2011)***). We performed initial feature selection using the Gradient Boosted Trees (XGBoost) Classifier after hyperparameter tuning by the Optuna framework (***Akiba et al. (2019)***) and the shap library (***Lundberg and Lee (2017)***). Features were ranked for importance using their SHAP (SHapley Additive exPlanations) values. Using a backward elimination method, the top 23 features were selected and used to train models.

### Model Training and Performance

We used a labelled dataset of known genes, out of which we have labelled 429 genes found in the Pain Genes Database (PGD) (***LaCroix-Fralish et al. (2007)***) and DOLORisk Priority Group (***Themistocleous et al. (2023)***) as Pain (P) and the remaining as Non-Pain (NP) genes (Figure 1B). These genes represent the gold-standard of highly confident targets in pain. Six classifiers, including random forest (RF), AdaBoost (Ada), gradient boost (GB), Gradient Boosted Trees (XGBoost); and Stacking and Voting ensembles were trained to classify pain/non-pain genes based on multi-omics, genomic and network topological data (Figure 2) (***Ho (1995); Schapire (2013); Friedman (2001); Chen and Guestrin (2016)***).

**Figure 2.**
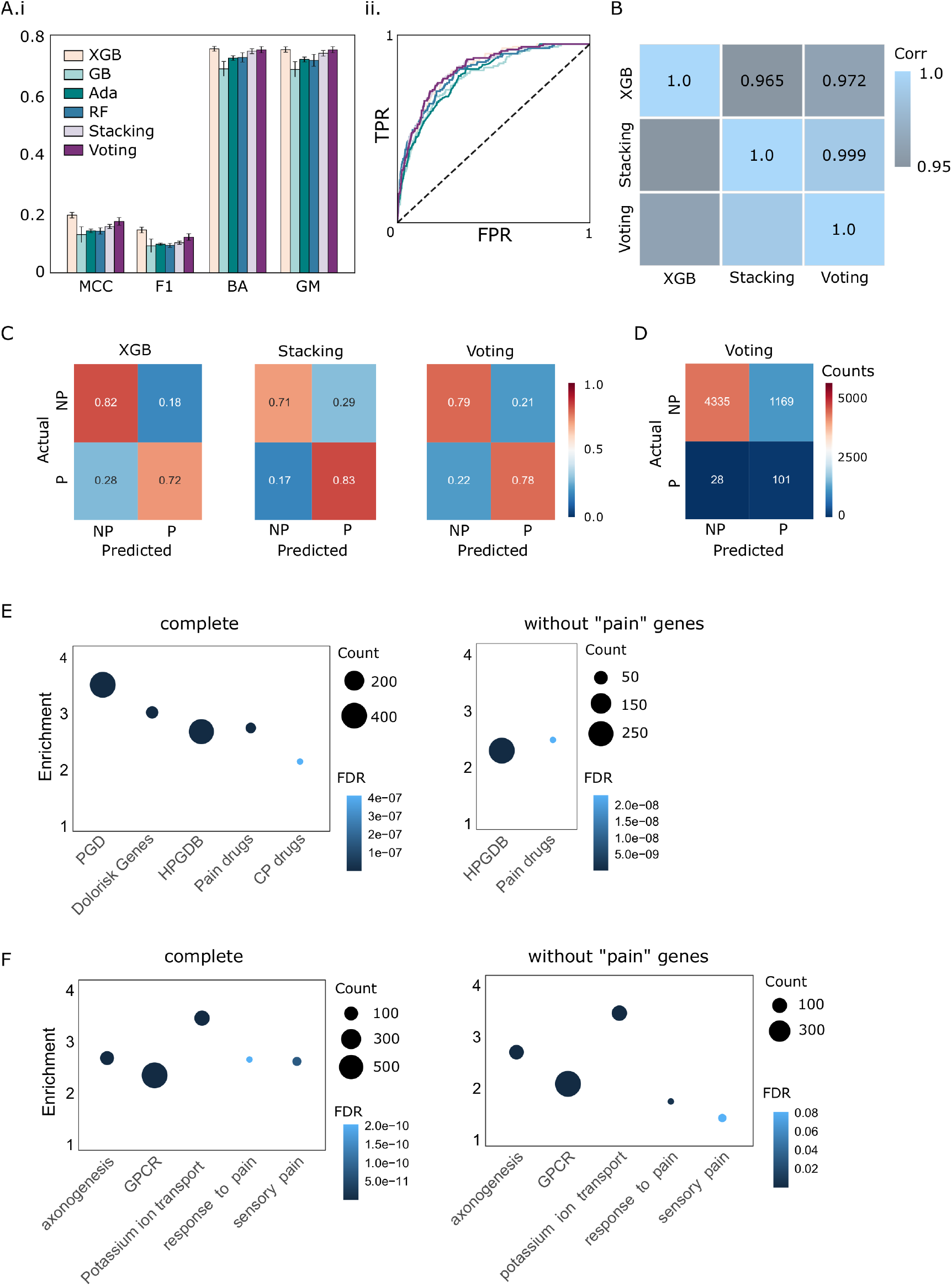
Classifier evaluation. A. Six classifiers were trained, with performance scores for (i) Mathew’s Correlation Coefficient (MCC), F1 score (F1), Balanced Accuracy (BA), and G mean (GM) evaluated. (ii) ROC curves show similar true positive rate (TPR) and false positive rate (FPR) across classifiers. B. Ranked prediction score correlations for the top three performing classifiers. C. Confusion matrix for the top three classifier in the test dataset. D. Confusion matrix by count for the top classifier in the test dataset, highlighting the imbalance in the dataset. E-F. Prediction scores for the “pain” class were ranked for enrichment analyses. Left; with all genes. Right; with all “no-pain” genes to prevent leakage. E. Predicted “pain gene” enrichment against curated known pain lists and approved drug targets. HPGBD: Human Pain Genetics Database, containing SNPs against relevant pain states/disorders. CP: chronic pain. F. Enrichment analyses against relevant biological pathways. Related to Supplemental Figure 1.

### Model Evaluation and Selection

We assessed model performance based on four metrics: F1 score, Matthews Correlation Coefficient (MCC), balanced accuracy (ACC), and geometric mean (GM) (Figure 2A-B) (***Chicco and Jurman (2020); Brodersen et al. (2010); Espíndola and Ebecken (2005)***). The voting classifier achieved the best performance (Voting; MCC = 0.1787 ± 0.0108, GM = 0.7581 ± 0.0132), followed by the XGBoost classifier (XGBoost; MCC = 0.1995 ± 0.0149, GM = 0.7535 ± 0.0192). The next best performance is the Stacking Classifier (Stacking; MCC = 0.1582 ± 0.0094, GM = 0.7452 ± 0.0132) and the Random Forest Classifier (RF; MCC = 0.1413 ± 0.0144, GM = 0.7183 ± 0.0266). Next, AdaBoost achieved a moderate performance (Ada; MCC = 0.1487 ± 0.0070, GM = 0.7300 ± 0.0061). Finally, GradientBoost Classifier achieved the worst performance (GB; MCC = 0.1325 ± 0.0171, GM = 0.6951 ± 0.0201). Results are summarized in Table 1.

**Table 1.** Model performance (mean ± SD) of six classifiers

| Algorithm | MCC | Balanced Accuracy | GM | F1 |
| --- | --- | --- | --- | --- |
| XGBoost | **$0.1995 \pm 0.0149$** | $0.7574 \pm 0.0169$ | $0.7535 \pm 0.0192$ | **$0.1530 \pm 0.0137$** |
| GradientBoost | $0.1325 \pm 0.0171$ | $0.6996 \pm 0.0155$ | $0.6951 \pm 0.0201$ | $0.0998 \pm 0.0165$ |
| AdaBoost | $0.1487 \pm 0.0070$ | $0.7331 \pm 0.0063$ | $0.7300 \pm 0.0061$ | $0.1022 \pm 0.0074$ |
| RandomForest | $0.1413 \pm 0.0144$ | $0.7290 \pm 0.0195$ | $0.7183 \pm 0.0266$ | $0.0934 \pm 0.0098$ |
| Stacking | $0.1582 \pm 0.0094$ | $0.7519 \pm 0.0115$ | $0.7452 \pm 0.0132$ | $0.1034 \pm 0.0072$ |
| Voting | $0.1787 \pm 0.0108$ | **$0.7586 \pm 0.0129$** | **$0.7581 \pm 0.0132$** | $0.1267 \pm 0.0091$ |

Here, we prioritized GM over MCC, while also looking at balanced accuracy and F1 scores. Together, these four metrics provide insight into the performance of imbalanced datasets, while GM controls accuracy in both classes and ranks higher classifiers that are equally good in both classes, regardless of their size (as it uses the ratios TPR, FPR). In our case, this is important as of course not many genes are validated as “pain genes”. This allows us to choose a classifier with a high number of true positives while still prioritizing the true negatives. (see Fig 2C).

The Voting Classifier ensemble was selected as the top performing model: Even so, we see a high correlation between prediction scores from top three highly ranked models (XGBoost, Voting Classifier, Stacking Classifier), suggesting that all three algorithms predict pain genes to a similar degree and that our predictions are sufficiently robust to not depend on a single classifier or set of parameters (Fig 2B). Although the voting classifier has a lower MCC and F1 score than XGBoost (2A), it predicts more true positive genes (Fig 2C-D), and has the highest GM and balanced accuracy, which are two important metrics in evaluating imbalanced datasets.

When building classiers, it is important to test the external validity of the model. In this design, we do not have a separate cohort to probe, as that would require a separate genome. Instead, we rely on relevant human genetics data, taking advantage of a curated list of single nucleotide polymorphisms (SNPs) relevant in pain from the Human Pain Genetics Database (HPGDB) ***Meloto et al. (2018)***. These represent relevant genetic polymorphisms in the context of pain across a broad range of conditions, in line with our original dataset. While some have been functionally validated and overlap with our gold-standard list of “pain genes”, many have not been followed up, representing likely, but unvalidated targets of pain. As such, these were not labelled as true pain genes in the original experimental design.

Functional GSEA was used to internally validate prediction scores from the top three highly ranked classifiers (XGBoost, Voting, and Stacking) using the known pain-related genes sets. In addition to internal validation against the sets used for labelling (Pain Genes Database and DOLORisk pain genes), prediction scores were externally validated by comparing them against the independent Human Pain Genetics Database as well as a curated list of pain-related drug-targets (Fig 2E). GSEAs were repeated for gene lists with true labels removed (that is, removing the 429 “pain” labelled genes where overlap occurs).

Crucially, external validation using genes in Human Pain Genetic Database after removing overlaps with our training data supports the use of the Voting Classifier, as it showed the highest enrichment for pain-related SNPs in the predicted pain ranking across genes in an unbiased dataset, as well as to drug targets for approved drugs relevant to pain and chronic pain (CP) (Figure 2E). GSEA plots for each classifier against the HPGDB without “pain” labels is shown in Supplemental Figure 1.

As expected, prediction results from the voting classifier also shows a high enrichment score for pain-related pathways including response to pain (GO:0048265) and sensory pain (GO:0051930) with and without the inclusion of pain-labelled genes (Fig. 2F).

### Feature analysis

In addition to studying which genes have the highest predictive pain scores, which features make up these predictions are also of interest. Because the voting classifier is an ensemble, SHAP values across classifiers were weighted and combined to reflect the most relevant features underlying the classification (Figure 3).

**Figure 3.**
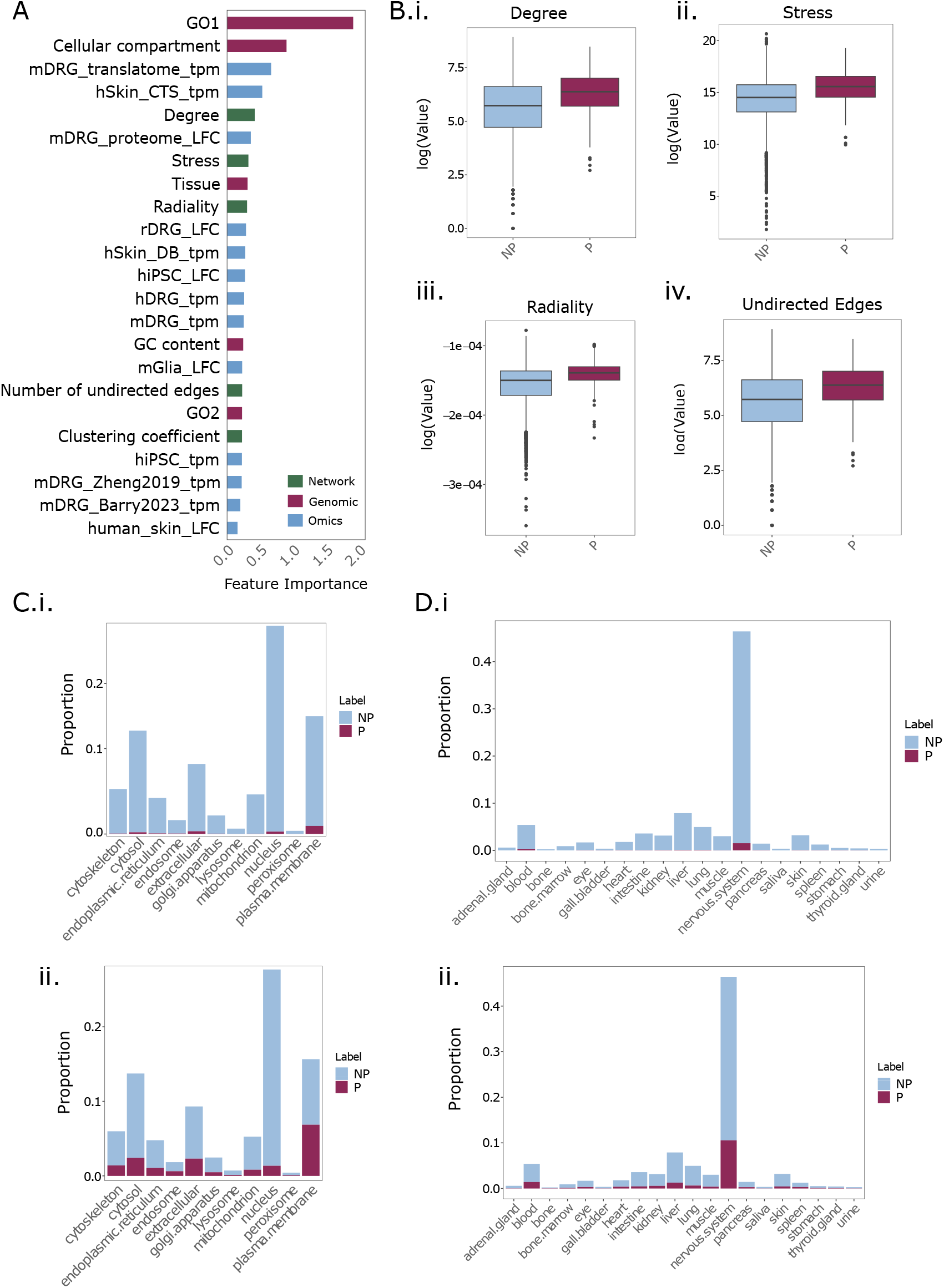
Voting classifier feature analysis. A. Ranked feature importance (weighted SHAP values) of the voting classifier. B. distribution of four top ranked network features in pain (P) and non-pain (NP) genes, extracted from the STRING DB through Cytoscape. ‘Degree’ refers to the number of edges linked to a node whereas ‘stress’ counts the number of shortest paths passing through a node. C. Proportion of genes in each cellular compartment (i) before and (ii) after prediction by the voting classifier. D. Proportion of genes in each tissue type (i) before and (ii) after prediction by the voting classifier. Species denoted as h (human), m (mouse), r (rat) (eg. hDRG_tpm = human DRG). TMP, transcripts per million; LFC, log fold change. Healthy human skin RNA-seq from two cohorts, CTS (carpal tunnel syndrome) and DB (diabetic) cohorts are highlighted as “hSkin_CTS_tpm” and “hSkin_DB_tpm” respectively.

The most highly informative feature was GO pathways. This is intuitive, given that well known pain genes used as true positive labels are associated with relevant GO pathways, thus classification by pathway is likely to be effective. Even so, we see here that even high order GO pathways from the GO slim collection are important, such that classification is not dependant on the smaller, pain-specific terms (e.g. “response to pain” and/or “sensory pain”, which are not present in the GO slim collection used to build the feature space)”.

The cellular component of the corresponding protein is the next highly ranked, with tissue expression following shortly there after. Notably, three network-based features extracted from the STRING database were also seen in the top 10 features (radiality, stress, and the number of undirected edges), suggesting that highly connected proteins are more likely to be involved in pain responses. These finding support a guilt-by-association approach showing that “pain” genes are likely to share similar functions as their interacting partners and aggregate in local interactome neighbours. Further exploration of network features shows that pain genes demonstrate higher radiality, stress, degree, and edge count (Fig. 3B). This suggest that they are likely to be hub proteins, which are the most highly connected central proteins in PPI networks (***Higurashi et al. (2008)***).

In terms of predictive “-omics” datasets, log fold changes in mDRG proteomics, as well as translatome murine expression studies also come up in the top 10, even above human DRG expression. Here, this may reflect that as the “functional building blocks” of a cell, changes in protein expression carries significant weight in predicting pain relevance. Alternatively, this may be a stronger comment towards a bias in the labels used, as some of our best documented “pain genes” were initially highlighted through rodent -omics studies, and/or that studies focus commonly focus on candidates with available antibodies due to technical limitations. As such, this also hints at the limitation of the true positive labels, which is discussed in below.

### Functional analyses

The predicted pain scores from the voting classifier were subjected to functional enrichment analysis.

#### GO and KEGG analysis

Given the high importance of GO in predicting pain-genes (Fig. 3A), we conducted GO functional and KEGG pathway enrichment analyses to explore which GO terms are enriched in pain genes (Fig 4A).

**Figure 4.**
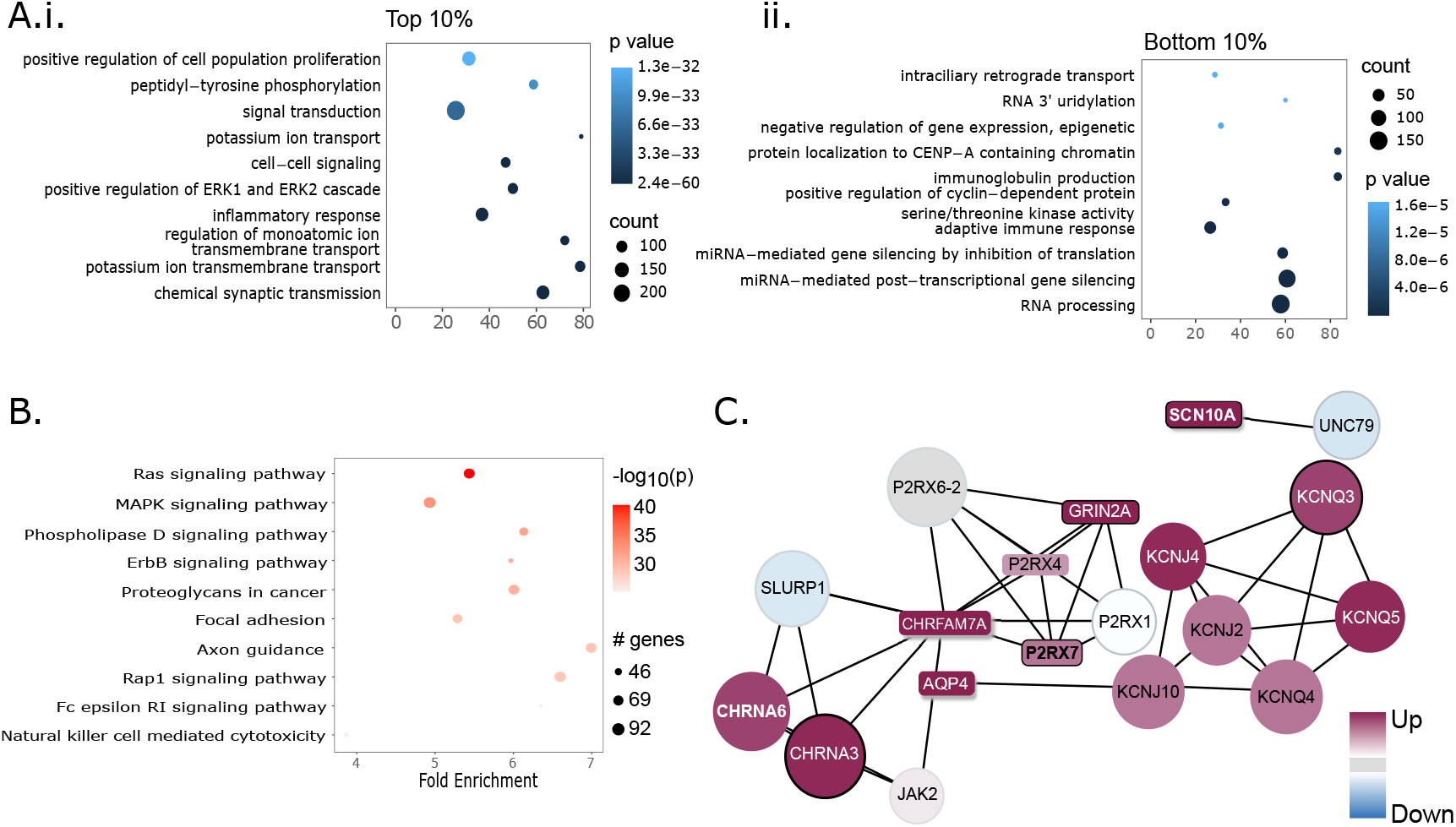
Functional analysis of high-ranked prediction scores from the Voting classifier. A. Top 10 GO term enrichment terms for top 10% and bottom 10% of ranked genes. B. Top 10 KEGG pathway terms for top 10% of ranked genes. C. PPI network of top 5 highly ranked genes by the Voting Classifier. Annotation from https://livedataoxford.shinyapps.io/drg-directory/: Colour gradient = Enrichment (red for PES > 0.02, blue for PES < 0.02); grey background = non-significant; Grey colour = absent from current dataset; Box shape = genes contained in the Pain Genes Database; Shadow = genes contained in the DOLORisk Priority Group; Black border = genes contained in the Human Pain Genetics Database.

GO term analysis of top 10% genes highlights GO term enrichment for potassium ion transport, regulation of membrane potential, positive regulation of cell proliferation, positive regulation of calcium ion concentration, positive regulation of ERK1 & ERK2 cascade, chemical synaptic transmission (Fig 4A.i). GO analysis of bottom 10% ranked genes show non-pain related GO terms, as expected (Fig 4A.ii).

KEGG analysis of top 10% genes shows up-regulation of Ras-, MAPK- and Erb-signalling pathways as promising targets for future study. It also involves the Natural Killer-mediated cytotoxicity pathway, suggesting this analysis can capture the critical roles of the immune system in pain, which was recently reviewed at ***Kim et al. (2023)***.

#### Network contextualization

Network features are highly ranked for predictive importance (Figure 3A-B). As such, we have integrated predictive scores of the voting classifier with the STRING DB through https://livedataoxford.shinyapps.io/drg-directory/ (“network analyses” tab, Figure 4C).

The network can be annotated using information on known pain-associated genes from several sources: DOLORisk Priority Group, Human Pain Genetics Database, and the Pain Genes Database (***Themistocleous et al. (2023); Meloto et al. (2018); LaCroix-Fralish et al. (2007))***. In addition, users can enrich these networks by using data from pain-focused gene expression studies to highlight genes that change expression in each condition or pairs of genes showing correlated expression patterns across different experiments.

The top 5 highly ranked genes are input to construct a PPI network (Fig 4C). Most of the extracted proteins demonstrate a high score, emphasising the significance of PPI networks in pinpointing disease-related proteins by elucidating their functional correlations with known pain genes.

## Discussion

The current study proposed a gene-centred machine learning approach to identify pain-related genes using multi-omics, PPI network and genomic data. ML has been used to identify disease biomarkers using multi-omics data (***Reel et al. (2021))***. However, due to the small sample sizes with high-dimensional features, training a large-scale generalizable ML model with multi-omics data alone can be challenging. Moreover, ML has also been successfully used to predict disease biomarkers using PPI network topological data (***Wu and Wang (2023)***). A study by Yu. et al used network topological features to predict proteins that may cause neurodegenerative disease, and used multi-omics data for validation and target selection (***Yu et al. (2020)***). Combining these approaches, we adopted a gene-centred approach to derive pain-related proteins from multi-omics, PPI networks, and genomic features. While previous studies focus more on the performance of the prediction algorithms, we brought attention to the biological explanation of predictive features and provided a list of genes as promising targets for future pain studies.

Out of six classifiers used, the voting classifier achieved the best performance (MCC = 0.1787 ± 0.0108, GM = 0.7581 ± 0.0132). It predicts the class label based on the argmax of the sums of the predicted probabilities by four individual classiers: XGBoost, AdaBoost, Stacking Classifier, and GradientBoost Classier.

Because many pain genes are known from rodent studies in the Pain Gene Database, external validation against the HPGDB and drug targets relevant to pain was crucial to establishing how to extrapolate probabilistic scores. The strong enrichment of the HPGDB, even after removing the gold-standard pain genes suggest these results can a) be extrapolated to humans, and b) are relevant in the context of human genetic studies. The enrichment of “pain drug” targets in high-ranking predictive scores further suggests that we are capturing a population of potentially-druggable targets (Figure 2E).

This study included data from multiple species, so we are likely capturing targets relevant cross-species, opposed to human-specific targets. Even so, gene “conservation scores” are themselves not predictive (Figure 3).

### Feature exploration

GO and KEGG functional analysis were conducted to identify enriched GO terms associated with pain genes. Analysis of the top 10% ranked genes revealed molecular functions closely linked to neuropathic pain pathogenesis. Among the enriched terms, five are linked to membrane transport processes (potassium ion transport, chemical synaptic transmission, regulation of membrane potential, regulation of ion transmembrane transport, and positive regulation of cytosolic calcium ion concentration), likely contributing to neuronal hyperexcitability associated with neuropathic pain (***Choi et al. (2023)***). The molecular functions of inflammatory response and cell-cell signalling are associated with production of pro-inflammatory molecules that sensitise nociceptive neurons underlying pain sensation (***Ji et al. (2016)***). The ERK1/2 is a characterised important signalling pathway in pain, and its activation is engaged in modulating the pain sensitivity (***Kondo and Shibuta (2020)***). GO analysis of bottom 10% ranked genes show non-pain related GO terms, as expected (Fig. 4A.ii).

KEGG analysis of the top 10% genes also revealed pathways associated with pain pathogenesis and progression. Ras signalling, implicated in NGF signalling through TrkA via the PI3K/Ras pathway leading to TRPV1 activation, is crucial in pain sensation (***Bonnington and McNaughton (2003)***). Ras is also frequently discussed in relation to the Ras/Raf/MAPK pathway: both pathways are enriched here. MAPK has a long-standing role in path pathophysiology ***Ji et al. (2009***), thus dissecting the interaction with Ras could lead to new avenues of druggable targets. Additionally, Rap1 signalling has also been described in the context of inflammatory pain through Epac1 (***Singhmar et al. (2016)***).

NRG1/ErbB signalling is significant in spinal cord injury (SCI)-induced chronic neuropathic pain (***Tao et al. (2013)***), and NRG1 has been implicated in axonal development as well as regeneration after nerve injury in the periphery ***Fricker et al. (2009***, 2011). Given the relevance to the neuropathic pain model of SCI, as well as recent work highlighting neuronal death after traumatic nerve injury in mice ***Cooper et al. (2024***), one can speculate how this pathway may be highly relevant across neuropathic pain conditions where axonal damage occurs. In line with this, the Natural Killer-mediated cytotoxicity pathway underscores cytotoxic immunity’s role in response to nerve injury (***Davies et al. (2019***, 2020)). Together, this suggests these are highly relevant candidates for further study.

Cellular compartment is the feature with second highest importance. Membrane proteins are heavily involved in pain reception. During nociception, high threshold stimuli that could lead to injury result in activation of ligand gated ion channels such as Transient Receptor Potential (TRP) channels in nociceptors in the periphery; subsequent opening of cation channels (potassium and sodium channels) results in depolarization and action potential propagation along afferent sensory fibres to the dorsal horn synapse (***Middleton et al. (2021)***). Consequently, the cellular compartment containing the highest number of pain-related genes is the plasma membrane (Fig. 3C). Interestingly, there are also a large number of pain-related genes in the nucleus, cytosol, and extracellular space, which serve as interesting insights for future target discovery.

While some features are highly predictive, others, such as species conservation rank less important in the current study. Here, it is unclear if this is a true trend, or if our bias in rodent studies covers a trend.

### Network Analysis

We developed and employed the Pain RNAseq Hub (https://livedataoxford.shinyapps.io/drg-directory/) to visualize the top 5 pain genes within their protein-protein interaction (PPI) network context, incorporating multi-omics data and pain-related annotations.

The extracted proteins include various characterised pain-related genes including CHRNA6, SCN10A, P2RX7, KCNQ5, KCNQ4, KCNQ3, CHRNA3, AQP4, JAK2, P2RX4. More importantly, we found several previously uncharacterised pain-related genes, which will be discussed below.

One such example is ADORA2A, a gene that encodes the adenosine A2A receptor. Binding of adenosine to the adenosine A2A receptor during stress initiates potentially destructive inflammatory cascades that lead to the activation of immune cells and release of proinflammatory mediators ***Flögel et al. (2012***). It was found that prolonged accumulated circulating adenosine contributes to chronic pain by promoting immune-neuronal interaction and revealed multiple therapeutic targets (***Hu et al. (2016)***). A2A receptor agonists have been shown to block adenosine and thus inhibit the release of proinflammatory mediators (***Cekic and Linden (2016)***) while related genes, ADORA2B and ADORA3, can cause nociceptor hyperexcitability and promote chronic pain (***Wahlman et al. (2018); Middleton et al. (2021)***). Together, this makes ADORA2A an attractive target to follow up.

Another interesting gene for future exploration is CHRFAM7A, a uniquely human fusion gene that functions as a dominant negative regulator of alpha 7 acetylcholine nicotinic receptors (*α*7nAChR). Recently, CHRFAM7A was found to contribute to exacerbating inflammation and tissue damage associated with osteoarthritis, and thus being a novel genetic risk factor and therapeutic target for pain (***Courties et al. (2023)***).

Lastly, UNC79 is an auxiliary subunit of the NALCN channel, which carries depolarizing sodium (Na+) leak currents to regulate the resting membrane potential of many neurons to modulate pain sensitivity (***Ren (2011)***). UNC79 and UNC80 are HEAT-repeat proteins that docks intracellularly onto the NALCN–FAM155A pore-forming subcomplex and are important for regulating the gating of NALCN. A recent study shows that the NALCN channel contributes to neuronal sensitization in neuropathic pain (***Zhang et al. (2021)***), and this result may lead experimental research to examine the regulatory mechanisms of NALCN by UNC79 and their associations with neuropathic pain in detail.

To encourage researchers from other fields to integrate multiple datasets, we provided a flexible, reproducible, and easy-to-understand code template (https://github.com/aliibarry/omics-database), as well as a tutorial, paving the way for better data utilisation and increased impact of individual -omics studies in biomarker discovery.

### Limitations

This research faces certain limitations, with one significant constraint being the limited number of pain-related genes that have been labelled, resulting in a class imbalance (P/NP = 1/40). Additionally, the relative difficulty in validating candidates poses a challenge in obtaining additional labels. To address these limitations, we employed multiple strategies: 1) utilising MCC and GM as the training metrics for model optimization, as they are more resilient to imbalance, and 2) incorporating ensemble classifiers into our approach.

### Future directions

As additional pain-related genes are discovered, it becomes possible to categorize different pain types, such as neuropathic, inflammatory, or cancer pain, which will ultimately refine gene prediction models specific to each subtype, enhancing accuracy. Building on this, the ability to probe tissue-specific mechanisms, opposed to a broad peripheral nervous system focus, will further enhance our knowledge. As new -omics datatypes continue to evolve and improve, such as epigenomics and human genetic studies, our ability to predict relevant candidates will also advance.

Furthermore, future research can focus on employing the existing machine learning framework to analyze gene expression patterns across a range of pathological conditions, such as neurodegenerative diseases, psychiatric disorders, and cancer, thereby broadening clinical impact and therapeutic understanding.

## Conclusions

This study uses large-scale multi-omics, PPI network, and genomic data to predict potential pain-related genes. Based on predicted pain scores, a number of hub proteins were selected as promising studies for future studies. A shiny app, accompanied by code template, is developed for further exploration of pain genes in the context of their PPI networks. Together, the findings and methodology presented in this study not only shed light on future directions in pain research, but also offer a valuable framework that can be adapted and applied to other fields for biomarker discovery.

## Methods and Materials

### Classification labels

The purpose of this study was two-fold: 1) explore which factors help predict if a gene is involved in pain, and 2) identify novel candidates for follow-up studies. To these effect, we used a rigorous labelling system to denote Pain (P) and Non-Pain (NP) genes, encompassing a variety of pain conditions. Here, we required high confidence levels in the true “pain” designation and assume that a subset of pain genes to be discovered within the list of “non-pain” genes.

Together, 429 “pain genes” from gold-standard, experimentally validated lists were used, including those from the Pain Genes Database (PGD) (***LaCroix-Fralish et al. (2007)***) and from the DOLORisk Priority Group (***Themistocleous et al. (2023)***) to label genes as Pain (P), the remaining known genes were labelled as Non-Pain (NP) genes. The PGD contains pain-related phenotypes (both acute and injury-induced) of transgenic mice, whereas the DOLORisk Priority Group includes genes shown to have a causal (Tier 1) role in human neuropathic pain based on casual variants in multiple, independent families. Tier 2 genes have been implicated in human neuropathic pain, but do not meet the criteria set out in Tier 1 and were also included as “pain genes”. Tier 3 genes, which as described as “of interest, as determined by expert consensus” but have little or no published data in the context of human neuropathic pain were omitted from the “pain” label (ie. labelled “non-pain”) but were used as a comparator for downstream analyses.

Genes from the Human Pain Genetics Database (HPGDB) (***Meloto et al. (2018)***), which contains pain-associated SNPs from human GWAS studies were labelled as NP: While SNPs against painful conditions are considered relevant to pain, there is a lack of experimental validation specifically highlighting a functional role. Future work may implicate these more strongly, but in this study they are labelled “NP” due to the lack of experimental validation unless there was data from the Pain Genes Database or DOLORisk priority genes suggesting that they were pain genes. These were instead used for external validation of the classification “pain score” in the form of an GSEA enrichment analysis (described below).

### Data input and pre-processing

Fifty-eight input features were selected from three categories: genomic features, experimental -omics datasets, and network topological coefficients, which were mapped from the rodent to human genome using biomaRt where necessary (Table 2 and 3).

**Table 2.** Genomic and Network Features

| Engineered Features | Description |
| --- | --- |
| *Genomic Features* | *Durinck et al. (2009, 2005)* |
| Cellular compartment | The cellular compartment by which the gene has the highest expression |
| GC | % GC content |
| Chromosome name | Name of chromosome |
| Conservation score | The conservation score of the gene; calculated as the total conservation score of each base divided by the number of DNA bases. |
| GO1 & GO2 | PCA components after vectorisation and PCA transformation of Gene Ontology terms of each gene; The GO terms are filtered such that only terms that appear in less than 20% of the genes are retained. |
| Tissue | The tissue with the highest expression |
| *Network Topological features* | *Gustavsen et al. (2019); Shannon et al. (2003)* |
| Average Shortest Path Length | Expected distance between two connected nodes |
| Betweenness Centrality | Control that this node exerts over the interactions of other nodes in the network |
| Closeness Centrality | How fast information spreads from a given node to other reachable nodes in the network |
| Clustering Coefficient | The number of triangles (3-loops) that pass through this node, relative to the maximum number of 3-loops that could pass through the node |
| Degree | The number of edges linked to a node |
| Eccentricity | The maximum non-infinite length of a shortest path between a node and another in the network |
| Neighborhood Connectivity | The connectivity of a node is the number of its neighbours. The neighbourhood connectivity of a node n is defined as the average connectivity of all neighbours of n |
| Undirected Edges | The number of undirected edges that are connected to a node |
| Radiality | A node centrality index |
| Stress | Counts the number of shortest paths passing through a node |
| Topological Coefficient | Relative measure for the extent to which a protein in the network shares interaction partners with other proteins |

**Table 3.** -omics features

| Engineered Features | Tissue | Species | Original Features | Description | Reference |
| --- | --- | --- | --- | --- | --- |
| Transcriptome DEG, mDRG | DRG | Mouse | LFC_CGRT_3D | LFC, subpopulations in mice after nerve injury | *Barry et al. (2023)* |
| Transcriptome DEG, mDRG | DRG | Mouse | LFC_CGRT_4W | LFC, subpopulations in mice after nerve injury | *Barry et al. (2023)* |
| Transcriptome DEG, mDRG | DRG | Mouse | LFC_CRTH_3D | LFC, subpopulations in mice after nerve injury | *Barry et al. (2023)* |
| Transcriptome DEG, mDRG | DRG | Mouse | LFC_CRTH_4W | LFC, subpopulations in mice after nerve injury | *Barry et al. (2023)* |
| Transcriptome DEG, mDRG | DRG | Mouse | LFC_MRTD_3D | LFC, subpopulations in mice after nerve injury | *Barry et al. (2023)* |
| Transcriptome DEG, mDRG | DRG | Mouse | LFC_MRTD_4W | LFC, subpopulations in mice after nerve injury | *Barry et al. (2023)* |
| Transcriptome DEG, mDRG | DRG | Mouse | LFC_TBAC_3D | LFC, subpopulations in mice after nerve injury | *Barry et al. (2023)* |
| Transcriptome DEG, mDRG | DRG | Mouse | LFC_TBAC_4W | LFC, subpopulations in mice after nerve injury | *Barry et al. (2023)* |
| Transcriptome DEG, mDRG | DRG | Mouse | LFC_TDNV_3D | LFC, subpopulations in mice after nerve injury | *Barry et al. (2023)* |
| Transcriptome DEG, mDRG | DRG | Mouse | LFC_TDNV_4W | LFC, subpopulations in mice after nerve injury | *Barry et al. (2023)* |
| Transcriptome DEG, mDRG | DRG | Mouse | LFC_balb | LFC values of mouse DRG gene expression after SNI | *Baskozos et al. (2019)* |
| Transcriptome DEG, mDRG | DRG | Mouse | LFC_b10d2 | LFC values of mouse DRG gene expression after SNI | *Baskozos et al. (2019)* |
| Transcriptome expression, mDRG subpopulations | DRG | Mouse | subtype_tpm | TPM counts of genes from transgenically-labelled subpopulations of neurons in male and female mice | *Barry et al. (2023)* |
| Transcriptome expression, hSkin (healthy) | Skin | Human | hSkin_DB_tpm | Naive gene expression counts in human skin | *Baskozos et al. (2022)* |
| Transcriptome DEG, hSkin from painful vs painless DB patients | Skin | Human | LFC_Diabetes | LFC values of human DPN patients vs control | *Baskozos et al. (2022)* |
| Transcriptome DEG, hSkin from painful vs painless DB patients | Skin | Human | LFC_Diabetes_male | LFC values of human DPN patients vs control, males | *Baskozos et al. (2022)* |
| Transcriptome DEG, hSkin from painful vs painless DB patients | Skin | Human | LFC_Diabetes_female | LFC values of human DPN patients vs control, females | *Baskozos et al. (2022)* |
| Transcriptome expression, hSkin (healthy) | Skin | Human | hSkin_CTS_tpm | Naive gene expression counts in human skin | *Baskozos et al. (2020)* |
| Transcriptome DEG, hSkin from carpal tunnel patients, pre- and post- surgery | Skin | Human | LFC_skin | LFC of the gene expression in skin | *Baskozos et al. (2020)* |
| Transcriptome expression, iPSC | iPSC | Human | iPSC_tpm | Gene expression counts of iPSC cells | *Clark et al. (2021)* |
| Transcriptome DEG, iPSC | iPSC | Human | LFC_iPSC_young | LFC values of iPSC cells following nerve injury, young | *Clark et al. (2021)* |
| Transcriptome DEG, iPSC | iPSC | Human | LFC_iPSC_old | LFC values of iPSC cells following nerve injury, old | *Clark et al. (2021)* |
| Transcriptome DEG, iPSC | iPSC | Human | LFC_iPSC | LFC values of iPSC cells following nerve injury | *Clark et al. (2021)* |
| Transcriptome expression, mDRG | DRG | Mouse | mDRG_sham_tpm | TPM, sham nerve injury DRG | *Baskozos et al. (2019)* |
| Transcriptome DEG, rat | DRG | Rat | SNI_rat_LFC | LFC, rat DRG model of SNI | *Maratou et al. (2008)* |
| Transcriptome DEG, rat | DRG | Rat | SNT_rat_LFC | LFC, rat DRG model of SNT | *Vega-Avelaira et al. (2009)* |
| Transcriptome DEG, rat | DRG | Rat | LFC_hiv | LFC, rat DRG model of HIV-associated neuropathic pain | *Maratou et al. (2008)* |
| Transcriptome DEG, rat | DRG | Rat | LFC_bone_cancer | LFC, rat DRG model of bone cancer | *Perkins et al. (2013)* |
| Transcriptome DEG, glial injury | DRG SG | Mouse | LFC_d3_glial | LFC, transcriptional fingerprint of satellite glial cells following peripheral nerve injury, day 3 | *Jager et al. (2020)* |
| Transcriptome DEG, glial injury | DRG SG | Mouse | LFC_d8_glial | LFC, transcriptional fingerprint of satellite glial cells following peripheral nerve injury, day 8 | *Jager et al. (2020)* |
| Transcriptome DEG, glial injury | DRG satellite glia | Mouse | LFC_d14_glial | LFC, transcriptional fingerprint of satellite glial cells following peripheral nerve injury, day 14 | *Jager et al. (2020)* |
| Translatome expression | DRG nociceptors | Mouse | trans_tpm | TPM, nociceptor translatomes data of in male and female mice following nerve injury | *Tavares-Ferreira et al. (2022)* |
| Translatome expression, normalized | DRG nociceptors | Mouse | transnorm_tpm | TPM, nociceptor translatomes data of in male and female mice following nerve injury, normalised (as published) | *Tavares-Ferreira et al. (2022)* |
| Proteome DEG (FDR), mSC | SC | Mouse | sc_lfc | LFC, protein expression in spinal cord after SNI in male mice | *Barry et al. (2018)* |
| Proteome DEG (LFC), mDRG | DRG | Mouse | drg_lfc | LFC, protein expression in DRG after SNI in male mice | *Barry et al. (2018)* |
| Proteome DEG (LFC), mSN | SN | Mouse | sn_lfc | LFC, protein expression in the sciatic nerve (SN) after SNI in male mice | *Barry et al. (2018)* |
| Proteome DEG (FDR), mSC | SC | Mouse | sc_padj | FDR, protein expression changes in the spinal cord (SC) after SNI in male mice | *Barry et al. (2018)* |
| Proteome DEG (FDR), mDRG | DRG | Mouse | drg_padj | FDR, protein expression changes in the dorsal root ganglia (DRG) after SNI in male mice | *Barry et al. (2018)* |
| Proteome DEG (FDR), mSN | SN | Mouse | sn_padj | FDR, protein expression changes in the sciatic nerve (SN) after SNI in male mice | ***Barry et al. (2018)*** |
| Transcriptome DEG, hiPSC-SN vs hiPSC | hiPSC-SN | Human | LFC_ipSC | LFC, gene expression in iPSC cells | ***McDermott et al. (2019)*** |
| Transcriptome expression, hDRG | DRG | Human | hdrg_tpm | TPM, naïve human DRG | ***Ray et al. (2022)*** |
| Transcriptome expression, mDRG subpopulation | DRG | Mouse | TPM_zheng | TPM, naïve mouse nociceptor | ***Zheng et al. (2019)*** |

#### Genomic features

The genomic features used in the model include the cellular compartment with the highest gene expression, the GC content (percentage of GC nucleotides) in the gene sequence, the chromosome name, and the conservation score of the gene indicating its evolutionary conservation. Conservation is calculated as the total conservation score of each base divided by the number of DNA bases. These features are retrieved using the biomaRt package (***Durinck et al. (2005)***) as well as the STRING DB plugin through Cytoscape, as discussed below. Both cellular compartment and tissue expression were extracted through the Cytoscape plug-in, representing the compartment and tissue in which the protein is the most highly expressed. These, along with the chromosome names were then vectorized using ‘OrdinalEncoder()’.

GO terms of each gene are also retrieved from Ensembl using ‘goslim_goa_accession’, with terms appearing in less than 20% of the genes retained. The GO “slim” dataset used provides an overview of biological function, retaining high order classifications without the specific terms such as “response to pain” which may affect classification through data leakage. Here, we verified that no terms containing the work “pain” were included during feature space generation. These terms were vectorised via term frequency - inverse document frequency (TF-IDF), which reduces the weight of frequent terms and increases the weight of rare ones. Dimensionality was reduced by PCA, where the data is projected in the subspace of a few principal components that explains most of the observed variance. With this, they can be visualised to assess the clustering of P vs NP genes.

#### Omics datasets

A curated selection of high quality -omics datasets were used to reflect diversity in species, relevant tissue, and high-throughput method. Together, this includes a mix of transcriptomic, translatomic, and proteomic datasets from mouse, rat, and human. Various tissues were included across the somatosensory pathway, both in the context of naïve expression as well as fold changes in injured states. This includes skin, nerve, dorsal root ganglia (DRG), and spinal cord, with a slight bias towards mouse DRG due to the sheer number of studies across pain conditions by the field. Where possible, human expression data and differential gene expression was included. A full list of datasets are available in Table 2 and 3.

#### Network Topological Features

The STRING DB was used to calculate network protein-protein interaction scores for human protein-coding genes (***Szklarczyk et al. (2021)***). Here, 11 general topological features were calculated using the software Cytoscape plugin NetworkAnalyzer using default parameters (Table 2 and 3). Features with zero variance (eg. PartnerOfMultiEdgedNodePairs and IsSingleNode) were not included. More detailed explanations and mathematical formulae can be found in the online help document of NetworkAnalyzer (https://med.bioinf.mpi-inf.mpg.de/netanalyzer/help/2.7/).

### Data pre-processing and Feature Engineering

Data processing was performed in Python, using Sklearn (***Pedregosa et al. (2011)***). Numerical features, including network topology, conservation scores, and -omics data were scaled using ***M**in**M**ax**S**caler*(*feature*_*r*_*ange* = (−1, 1)). Categorical features were converted as described above prior to scaling with using ***M**in**M**ax**S**caler*(*feature*_*r*_*ange* = (−1, 1)).

For pairs of highly correlative features (correlation coefficients > 0.75), the feature with smaller variance was removed. Including highly correlated features can lead to overfitting and decrease the model’s interpretability.

The data was split into training (70%) and validation (30%) sets, stratified by label. Numerical features were centred and normalised separately for the training and validation set using min-max normalisation from the Sklearn library. This step ensures unbiased feature comparisons and improves the stability and convergence of ML algorithms.

To improve the model’s effectiveness by reducing complexity, we calculated composite LFC values for bulk transcriptomics for every tissue and species. This involved categorizing datasets according to tissue and species. Within each group, we determined the mean LFC value for every gene, after setting non-significant entries (FDR > 0.05) to zero.

### Evaluation Metrics

This dataset is featured by a severe class imbalance due to our stringent labelling of true “pain” genes. To tackle the class imbalance, four metrics (Matthews Correlation Coefficient (MCC, 1), geometric mean (GM, 2), F1 score (F1, 3), and balanced accuracy (BA, 4)) were chosen to be maximised during model selection and for benchmarking the best performing models during validation. These metrics have been shown to be robust in class imbalances. Equations for each metric are provided, with TP = true positive, TN = true negative, FP = false positive, FN = false negative.

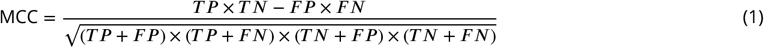

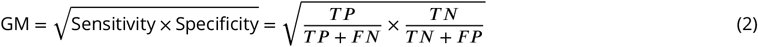

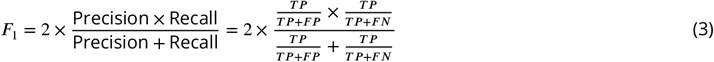

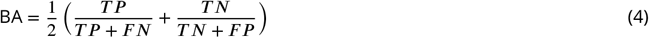

### Hyperparameter Tuning & Feature Selection

Initial feature selection was performed using the XGBoost Classifier after hyperparameter tuning by Optuna framework (***Akiba et al. (2019)***) and the shap library (***Lundberg and Lee (2017)***). The optimal number of features was determined through backward elimination, removing one feature at a time. The combinations of features with the greatest GM and the greatest MCC score were then used to train models. The final 23 features are highlighted in Figure 3A.

Features were ranked for importance using their SHAP (SHapley Additive exPlanations) values. These values are derived from cooperative game theory and represent the importance of each feature to a given model. They can be approximated using ‘shap.TreeExplainer()’ in Python by extracting the ‘shap_value’ from the test data.

### Model Training

Six machine learning models, including RandomForest Classifier, AdaBoostClassifier, GradientBoostingClassifier, XGBoost Classifier, Stacking Classifier, and Voting Classifier were used for classifying pain/non-pain genes (***Ho (1995); Schapire (2013); Friedman (2001); Chen and Guestrin (2016)***). After using the backward elimination method, optimizing for GM and MCC, 23 features were used (Figure 3A). The Stacking Classifier (final_estimater = ‘logistic regression’) and Voting Classifier(voting = ‘soft’) consists in stacking the output of RandomForest Classifier, AdaBoostClassifier, GradientBoostingClassifier, and XG-Boost Classifier.

Hyperparameter tuning was conducted to optimise the summed GM, MCC, BA, and F1 scores of models using the Optuna framework ***Akiba et al. (2019***). The weights of the four individual classifiers in the voting classifier were tuned using gridsearch. During the training step, a 10-fold cross-validation was utilised to assess the models’ performance. The full list of parameters are available in a Jupyter Notebook at https://github.com/aliibarry/omics-classifier/.

The voting classifier is a weighted ensemble of the base classifiers. A permutation function generates all possible orderings of the list, and the weight is selected by calculating and selecting the highest GM score of the classifier with all permutations of weights. Here, the weights [4,2,1,3] correspond to the estimators [xgbm, gb, ada, rf]. We set the voting to ‘soft’, by which the classifier predicts the class label based on the argmax of the sums of the predicted probabilities by each individual classifier. The class probabilities predicted by each classifier are multiplied by the weight before averaging (soft voting).

#### Model Validation

The gene ranking is derived based on the class probability of the best performing classifier. Internal validation of gene ranking is achieved by GSEA enrichment against the Pain Genes Database, which are derived from results of pain-relevant knockout studies, and the DOLORisk pain Genes. External validation is achieved by GSEA enrichment against Human Pain Genetics Database (HPGDB), which contains pain-associated genes from human GWAS studies (***Meloto et al. (2018)***). This was run with and without overlapping “pain” labelled genes to prevent any leakage from the original data, as some SNPs from the HPGDB have been validated functionally and are thus contained in our “pain gene” list. Data were also compared to drugs relevant to “pain”, “neuropathic pain”, and “chronic pain”, extracted from opentargets.org (***Ochoa et al. (2023)***). Here, only approved drugs were used for GSEA analyses.

#### Functional Analysis

Gene Ontology (GO) analysis was conducted on the top and bottom 10% ranked genes respectively using the goseq package in R to identify enriched GO terms associated with top ranking pain genes, using the complete set of GO terms (***Young et al. (2010)***). A significance threshold of adjusted p-value (Benjamini-Hochberg corrected) < 0.05 was applied to determine significantly enriched GO terms. KEGG analysis was done on the top 10% ranked genes using the pathfindR package (***Ulgen et al. (2019)***). Furthermore, the top 10% genes were subjected to GSEA against the HSPGB, pain-related GO terms (GO: 0051930, GO: 0071805, GO:0007409, GO:0048265, GO:0007186) and pain-unrelated GO terms (GO:0006357 and GO:0006355) using the clusterProfiler R package (***Wu et al. (2021)***).

## Data and code availability

No new datasets were generated from this study. Predicted pain scores and searchable datasets are searchable at https://livedataoxford.shinyapps.io/drg-directory/. A Jupyter Notebook for building the classifiers is available at https://github.com/aliibarry/omics-classifier. Database code, with a simplified example are available at https://github.com/aliibarry/omics-database. A manual for database development is available at https://aliibarry.github.io/database-book/.

## Author contributions

AMB, NZ, and GB designed the study with input from DB. NZ trained classifiers with input from AMB and GB. AMB and NZ developed the database and associated code/manual to improve data access with input from GB and DB. AMB and NZ drafted the manuscript and figures that were reviewed and approved by all authors.

## Acknowledgments

This work was funded in part by a Wellcome Investigator Grant to DB (223149/Z/21/Z), as well as the MRC (MR/T020113/1), and with funding from the MRC and Versus Arthritis to the PAINSTORM consortium as part of the Advanced Pain Discovery Platform (MR/W002388/1). GB is funded by Diabetes UK, grant number 19/0005984, MRC and Versus Arthritis through the PAINSTORM consortium as part of the Advanced Pain Discovery Platform (MR/W002388/1) and by the Wellcome Trust (223149/Z/21/Z). AMB is funded by the MRC and Versus Arthritis through the PAINSTORM consortium as part of the Advanced Pain Discovery Platform (MR/W002388/1). ChatGTP was used to help format equations in LaTeX for submission.

This research was funded in part by the Wellcome Trust [223149/Z/21/Z]. For the purpose of open access, the author has applied a CC BY public copyright license to any Author Accepted Manuscript version arising from this submission.

---

**Figure supplement 1.**
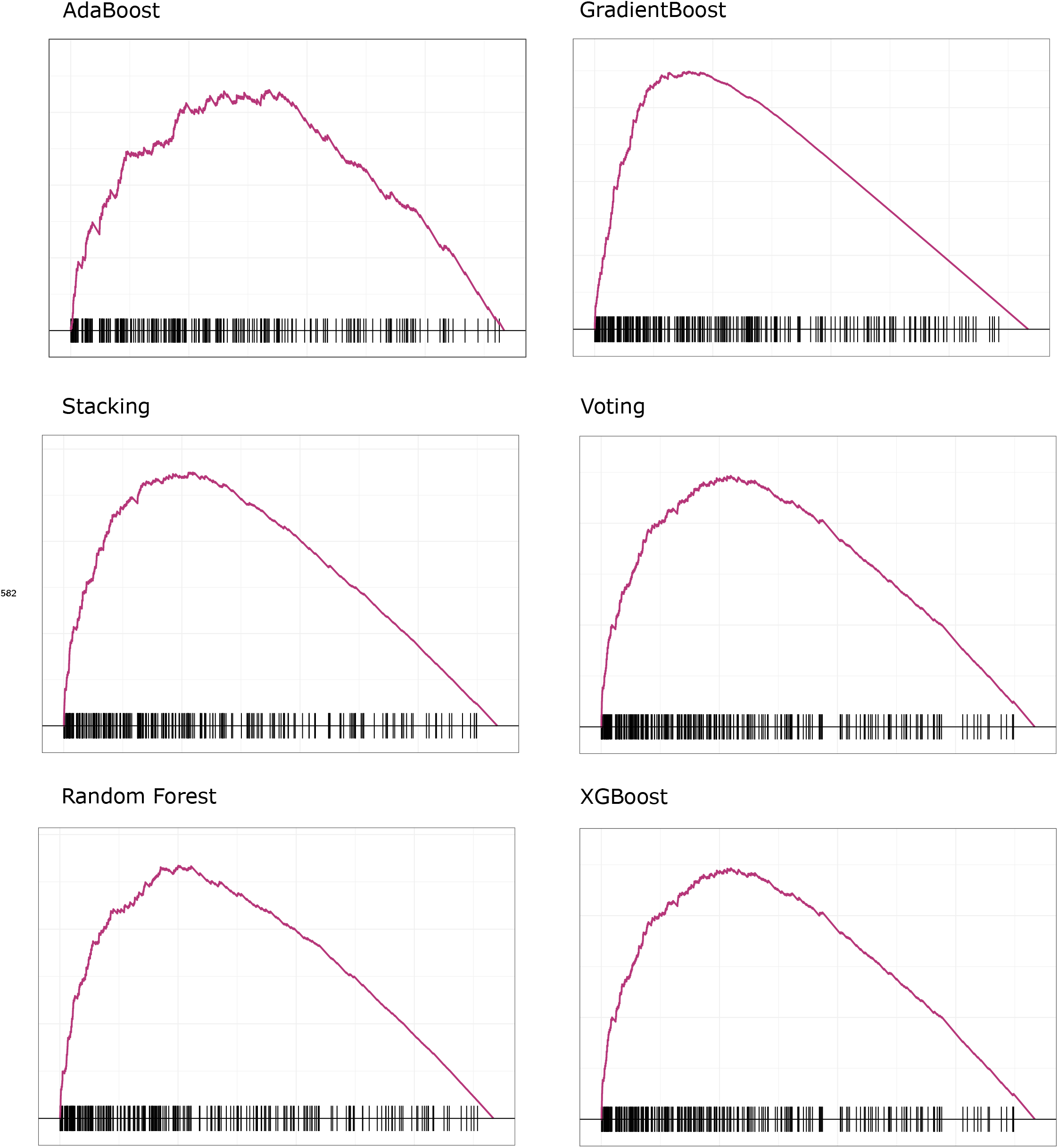
GSEA validation analyses against HPGDB without “pain” labels

